# Identification of conserved epitopes in SARS-CoV-2 spike and nucleocapsid protein

**DOI:** 10.1101/2020.05.14.095133

**Authors:** Sergio Forcelloni, Anna Benedetti, Maddalena Dilucca, Andrea Giansanti

**Affiliations:** Sapienza University of Rome, Department of Physics, P.le A. Moro 5, 00185 Rome, Italy; Sapienza University of Rome, Department AHFMO, Via A. Scarpa 14, 00161 Rome, Italy; Istituto Nazionale di Fisica Nucleare, INFN, Roma1 section 00185, Roma, Italy

**Author notes:** Corresponding Author: Sergio Forcelloni, Sapienza University of Rome, Department of Physics, P.le A. Moro 5, 00185 Roma, Italy.

**Keywords:** SARS-CoV-2, spike glycoprotein, nucleocapsid protein, conserved epitopes, conservation score, order-disorder propensity

## Abstract

Severe acute respiratory syndrome coronavirus 2 (SARS-CoV-2), which first occurred in Wuhan (China) in December 2019, is a novel virus that causes a severe acute respiratory disease. The virus spike glycoproteins and nucleocapsid proteins are the main targets for the development of vaccines and antiviral drugs, to control the disease spread. We herein study the structural order-disorder propensity and the rates of evolution of these two proteins to characterize their B cell and T cell epitopes, previously suggested to contribute to immune response caused by SARS-CoV-2 infections. We first analyzed the rates of evolution along the sequences of spike and nucleocapsid proteins in relation to the spatial locations of their epitopes. For this purpose, we compared orthologs from seven human coronaviruses: SARS-CoV-2, SARS-CoV, MERS-CoV, HCoV-229E, HCoV-OC43, HCoV-NL63, and HCoV-HKU1. We then focus on the local, structural order-disorder propensities of the protein regions where the SARS-CoV-2 epitopes are located. We show that the vast majority of nucleocapsid protein epitopes overlap the RNA-binding and dimerization domains and some of them are characterized by low rates of evolutions. Similarly, spike protein epitopes are preferentially located in regions that are predicted to be ordered and well-conserved, in correspondence of the heptad repeats 1 and 2. Interestingly, both the receptor-binding motif to ACE2 and the fusion peptide of spike protein are characterized by high rates of evolution, probably to overcome host immunity. In conclusion, our results provide evidence for conserved epitopes that may help to develop long-lasting, broad-spectrum SARS-CoV-2 vaccines.

## 1. Introduction

Coronaviruses are a large family of viruses which may cause illness in animals. In humans, seven coronaviruses are known to cause respiratory infections, ranging from the common cold to more severe diseases. The first two coronaviruses, human CoV-229E (HCoV-229E) and human CoV-OC43 (HCoV-OC43), were discovered in the 1960s, and cause relatively mild respiratory symptoms. Human severe acute respiratory syndrome coronavirus (SARSr-CoV) was identified in 2003, and causes flu-like symptoms and atypical pneumonia in the worst cases [Fouchier et al. 2003]. The human coronavirus NL63 (HCoV-NL63), identified in 2004, and the human CoV-HKU1 (HCoV-HKU1), described in 2005 [Woo et al. 2005], generally cause upper respiratory disease in humans, which may progress in lower respiratory infections [Van der Hoek et al. 2004]. More recently, the pathogenic Middle East respiratory syndrome (MERS-CoV) coronavirus, which appeared for the first time in 2012, was identified as the sixth human coronavirus [Zaki et al. 2012]. Finally, a previously unknown coronavirus probably originated in baths, SARS-CoV-2, was identified in December 2019 in Wuhan, China [Andersen et al. 2020,Gorbalenya et al. 2020]. SARS-CoV-2 caused an ongoing pandemic of severe pneumonia named coronavirus disease 19 (COVID-19), which has affected over 4 million people worldwide, and caused more than 300.000 deaths as May 13, 2020.

Currently, no vaccine for any of the known coronaviruses has been approved. Coronavirus vaccines can be live attenuated/ inactivated viruses, protein-based vaccines, DNA and mRNA vaccines [Zhang et al. 2020]. The viral spike (S) glycoprotein and the nucleocapsid (N) protein are two of the main targets for antibody production and for the development of vaccines and antiviral drugs [Zhang et al. 2020], due to their ability to trigger a dominant and long-lasting immune response.

The Spike protein is a large type I transmembrane protein of approximately 1400 amino acids. S protein is an attractive target for vaccine development, as its surface expression renders it a direct target for the host immune response. Spike proteins assemble into trimers on the virion surface to form the distinctive crown-like structure, and mediate the contact with the host cell by binding to ACE2 receptor, a process necessary for the virus entry. S protein contains 2 subunits: S1N-terminal domain, responsible for receptor binding, and S2 C-terminal domain, responsible for the fusion. The S2 subunit is the most conserved one, while the S1 subunit differs even within species of the same coronaviruses. The S1 contains two sub-domains (N-terminal and C-terminal), which are both show receptor-binding functions. The S2 domain contains two heptad repeats with hydrophobic residues, responsible for the formation of an α-helical coiled-coil structure that participate in the virus-host cell membrane fusion [Walls et al. 2020].

The Nucleocapsid protein regulates the viral genome transcription, replication and packaging, and it is essential for viability. The N protein is of potential interest for vaccine development as it is highly immunogenic and its amino acid sequence is highly conserved. This protein contains two structural domains: the N-terminal domain, that acts as a putative RNA binding domain, and a C-terminal domain, which acts as a dimerization domain [Tilocca et al. 2020].

T cell-responses against S and N proteins have been shown to be the most immunogenic and long lasting in SARS-CoV patients. Furthermore, B-cell antibody response against S and N proteins was also reported to be effective, although short-lived compared to the T cell-response. The search of T-cell and B-cell epitopes, which can stimulate a specific immune response against S and N proteins, represents a valuable strategy to identify targets for the development of a SARS-CoV-2 vaccine [Ahmed et al. 2020].

In a previous study, we showed that genes encoding N and S proteins tend to evolve faster than genes encoding matrix and envelope proteins [Dilucca et al. 2020]. This result suggested that the higher divergence observed for these two genes could represent a significant barrier in the development of antiviral therapeutics against SARS-CoV-2. Here, we perform a comprehensive analysis of the position-specific rates of evolution and the order-disorder propensities of the spike glycoprotein (S) and the nucleocapsid protein (N) of SARS-CoV-2. Thus, we provide an *in-silico* survey of the major nucleocapsid protein and spike protein epitopes and identify a subset of them that are well-conserved among homologues of human coronaviruses that could help design broad-spectrum vaccines against SARS-CoV-2.

## 2. Materials and Methods

### 2.1 Data sources

The complete coding genomic sequences of SARS-CoV-2 was obtained from NCBI viral databases, accessed as of 10May2020. In this study, we also considered six human coronaviruses: human CoV-229E (HCoV-229E), human CoV-OC43 (HCoV-OC43),human severe acute respiratory syndrome coronavirus (SARSr-CoV), human coronavirus NL63 (HCoV-NL63), human CoV-HKU1 (HCoV-HKU1), Middle East respiratory syndrome coronavirus (MERS-CoV), and the severe acute respiratory syndrome-related coronavirus 2 (SARS-CoV-2). We downloaded the coding sequences of these coronaviruses from the National Center for Biotechnological Information (NCBI) (available at https://www.ncbi.nlm.nih.gov/). For each virus, we investigated the evolutionary conservation and the structural disorder tendency of protein N (UniProt ID: P0DTC9) and S (UniProt ID: P0DTC2), which are regarded as important targets for the development of vaccines and antiviral drugs.

### 2.2 Sequence alignment

To explore the evolutionary relationships among the proteins N and S in the seven human coronaviruses here considered, the selected protein sequences were aligned by using Clustal Omega (https://www.ebi.ac.uk/Tools/msa/clustalo/) [Madeira et al. 2019]. This tool is a new multiple sequence alignment (MSA) program that uses seeded guide trees and HMM profile-profile techniques to generate alignments and phylogenetic tree of divergent sequences.

### 2.3 Disorder prediction

The structural order–disorder propensity of each protein was predicted by usingIUPred2A (https://iupred2a.elte.hu)[Mészáros et al. 2018], using the option for long disordered regions. Briefly, IUPred2A is a fast, robust, sequence-only predictor based on an energy estimation approach that allows to identify disordered protein regions. The key component of the calculations is the energy estimation matrix, a 20 by 20 matrix, whose elements characterize the general preference of each pair of amino acids to be in contact as derived from globular proteins. This prediction associates a score to each residue in the protein sequence ranging from 0 (strong propensity for an ordered structure) to 1 (strong propensity for a disordered structure), with 0.5 as the cut-off to classify residues as either ordered or disordered. In line with the original protocol [Dosztànyi et al. 2005], the position-specific estimations of the structural order–disorder propensity of each residue were then averaged over a window of 21 residues, and the average value was assigned to the central residue of the window (taking into account the limitations on both sides of the protein sequence).

### 2.4 Rate of evolution for site

We calculated the rate of evolution per site of the SARS-CoV-2proteins N and S relative to their orthologs in other six human coronaviruses using Rate4site (https://m.tau.ac.il/~{}itaymay/cp/rate4site.html) [Pupko et al. 2002].Rate4Site calculates the evolutionary rate at each site in the MSA using a probabilistic-based evolutionary model. This allows taking into account the stochastic process underlying sequence evolution within protein families and the phylogenetic tree of the proteins in the family. The conservation score at each site in the MSA corresponds to the site’s evolutionary rate. The position-specific estimations of the rate of evolution of each residue were then averaged over a window of 21 residues, and the average value was assigned to the central residue of the window (taking into account the limitations on both sides of the protein sequence). The size of the window was taken equal to that used above in section 2.3.

## 3. Results

### 3.1 Identification of conserved epitopes in spike and nucleocapsid proteins

The aminoacidic sequences of the nucleocapsid (N) protein and spike (S) glycoprotein from the seven human coronaviruses here considered (SARS-CoV-2, SARS-CoV, MERS-CoV, HCoV-229E, HCoV-OC43, HCoV-NL63, and HCoV-HKU1) were compared to assess the position-specific rates of evolution of these two proteins. We then investigated the relationships between the position-specific rates of evolution and the distribution of the epitopes that have previously been suggested to contribute to immune response caused by human SARS-CoV-2 infections. For this purpose, we considered the SARS-CoV-2B cell and T cell epitopes derived from N and S proteins by Ahmed et al. [Ahmed et al. 2020]. It is worth noting that these epitopes were also considered in another study aiming to provide a molecular structural rationale of the major nucleocapsid protein epitopes for a potential role in conferring protection from SARS-CoV-2 infection [Tilocca et al. 2020].We herein focused on the SARS-CoV-2 N and S proteins because both these two proteins are the main targets of vaccines and antiviral drugs due to their dominant and long-lasting immune response previously reported against SARS-CoV [Ahmed et al. 2020, Zhang et al. 2020].We aligned the homologous protein sequences in the seven human coronaviruses and used the resulting alignment to calculate a conservation profile by using the software Rate4site [Pupko et al. 2002] (see Materials and Methods). In Fig. 1, we report the profile obtained for the protein N, together with the functional regions/domains and the SARS-CoV-2 derived B cell and T cell epitopes.

**Figure 1:**
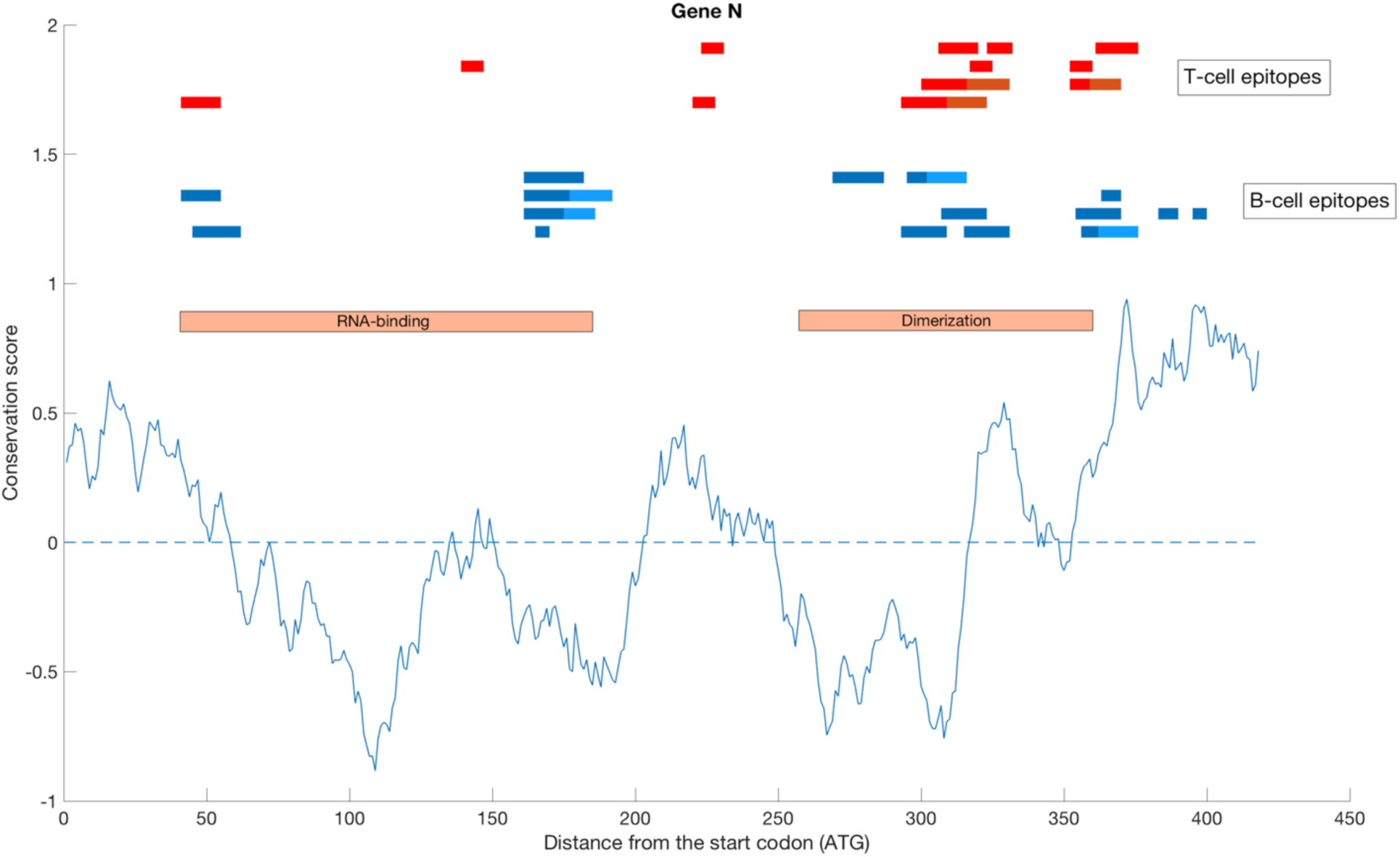
Conservation profile of SARS-CoV-2 protein N. The solid blue line represent the position-specific estimations of the rate of evolution of each residue averaged over a window of 21 residues around that position (taking into account the limitations on both sides of the protein sequence) as a function of the distance from the start codon. Thin horizontal dotted line represent the threshold value, above which the score is characteristic of disorder (0 for Rate4site). On the bottom, with two orange rectangles we show the RNA-binding domain and the dimerization domain. On the top, we report the SARS-CoV-2 derived B cell epitopes (in blue) and T cell epitopes (in red) by Ahmed et al. [Ahmed et al. 2020].

In this profile, values greater or less than zero reflect a faster or a slower evolution, respectively. We note that both the RNA-binding domain (region 41-186) and the dimerization domain (region 258-361) correspond to regions with values less than zero, implying a higher conservation with respect to the rest of the sequence. Although the general trend of the conservation profile hovers around 0, there are some regions where the conservation score is greater than one, indicating that these regions are likely to be under positive selection. Interestingly, the location of some of these regions corresponds to the presence of B cell and T cell epitopes (see Fig. S1 for the specific location and the sequences of each epitope). These segments could display high amino acid variability because amino acid diversity in these regions allows the virus to evade the host immune system recognition. Thus, we can conclude that the source of variability in this regions is likely to be the host immune response. However, it is worth noting the presence of epitopes in highly conserved regions spanning residues 150-200 and 250-315, which may potentially offer long-lasting protection against SARS-CoV-2.

Similarly, we study the conservation profile of the protein S and the distribution of the associated epitopes (Fig. 2).

**Figure2:**
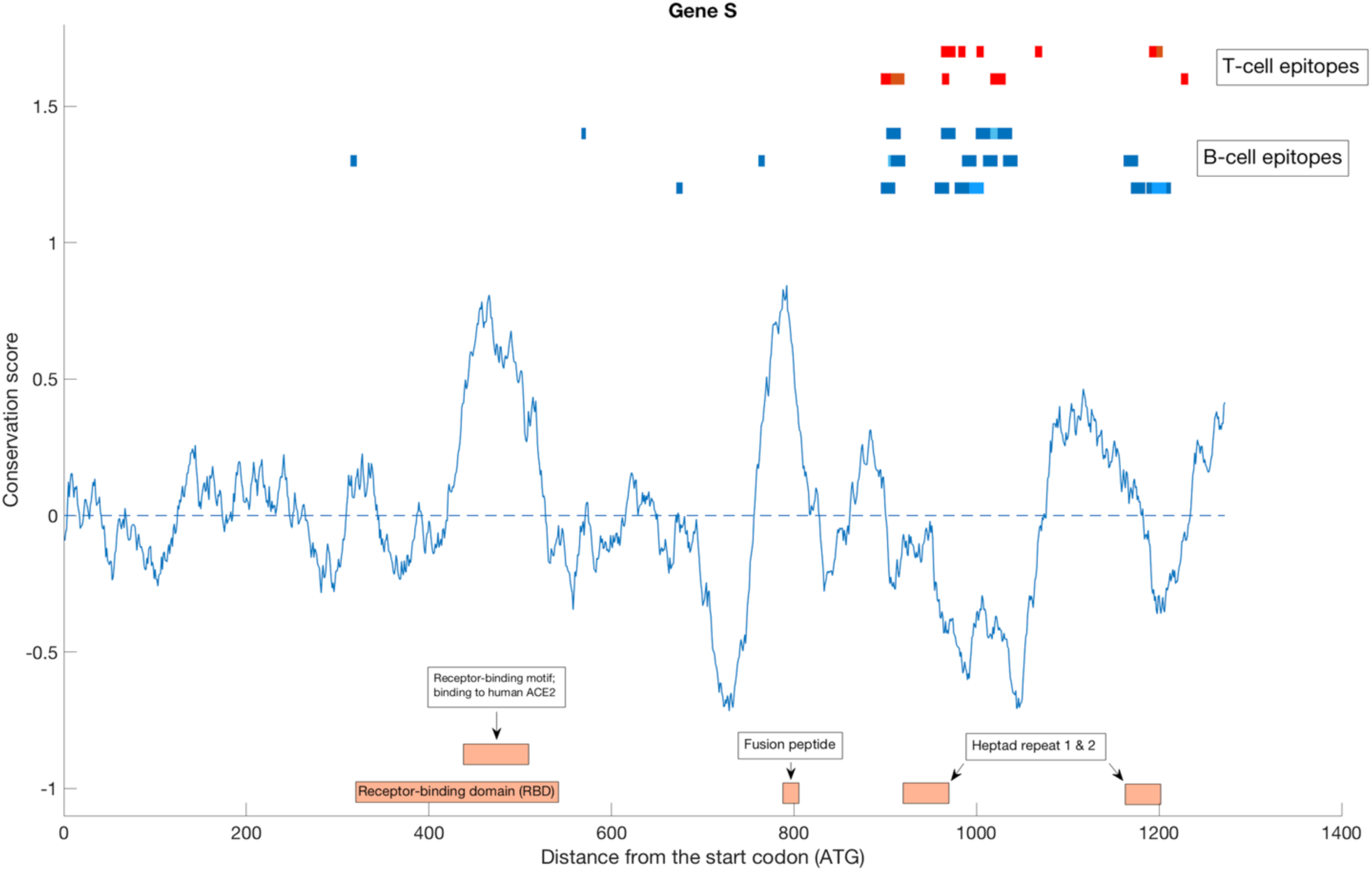
Conservation profile of SARS-CoV-2 protein S. The solid blue line represent the position-specific estimations of the rate of evolution of each residue averaged over a window of 21 residues around that position (taking into account the limitations on both sides of the protein sequence) as a function of the distance from the start codon. Thin horizontal dotted line represent the threshold value, above which the score is characteristic of disorder (0 for Rate4site). On the bottom, with orange rectangles we show the receptor-binding domain and its receptor binding motif to human ACE2, the fusion peptide, and the two heptad repeats. On the top, we report the SARS-CoV-2 derived B cell epitopes (in blue) and T cell epitopes (in red) by Ahmed et al. [Ahmed et al. 2020].

On the bottom of Fig. 2, we report the functional regions of protein S. The receptor binding domain (region 319-541) contains the receptor-binding motif to the human angiotensin-converting enzyme 2 (ACE2), which is an enzyme attached to the outer surface (cell membrane) of cells in the lungs, arteries, heart, kidney, and intestines, and it has been identified as the functional receptor for SARS-CoV-2 [Hamming et al. 2004]. As a transmembrane protein, ACE2 serves as the main entry point into cells for some coronaviruses, including HCoV-NL63, SARS-CoV, and SARS-CoV-2 [Fehr et al 2015; Li 2013; Kuba et al. 2005; Zhou et al 2020; Xu et al. 2020]. More specifically, the binding of the spike S protein of SARS-CoV and SARS-CoV-2 to the enzymatic domain of ACE2 on the surface of cells results in endocytosis and translocation of both the virus and the enzyme into endosomes located within cells [Wang et al. 2008; Millet and Whittaker 2018]. Interestingly, both the receptor-binding motif to ACE2 (region 437-508) and the fusion peptide (amino acids 788-806IYKTPPIKDFGGFNFSQIL for SARS-CoV-2), the segment of the fusion protein that inserts to a target lipid bilayer and triggers virus-cell membrane fusion, are characterized by high rates of evolution. Conversely, heptad repeats 1 and 2,which are known to play a crucial role in membrane fusion and viral entry [Liu et al 2004], show lower rates of evolution.

Moreover, we note that a large percentage of both B cell and T cell epitopes are located in the C-terminal region of Spike protein in correspondence of the heptad repeats 1 and 2 in the S2 domain (see Fig. S2 for details about the sequences of epitopes and their location along the sequence). Thus, at variance with the protein N, we note that spike protein epitopes are mainly located in the protein regions that are characterized by lower rates of evolution. This observation suggests that the immune system has adapted to recognize slowly evolving regions of the S protein.

### 3.3 Order–disorder propensities of proteins S and N and their associated epitopes

We predicted the structural order-disorder propensity proteins S and N to investigate the relationship between disordered structure and the spatial distribution of the SARS-CoV-2 derived B cell and T cell epitopes. To estimate the structural stability of a protein from its sequence without relying on the structure, we used the energy estimation approach at the core of the IUPred2A disorder prediction method (see Materials and Methods). In Fig. 3, we show the order-disorder propensity profile for protein N, together with the SARS-CoV-2 derived B cell and T cell epitopes.

**Figure 3:**
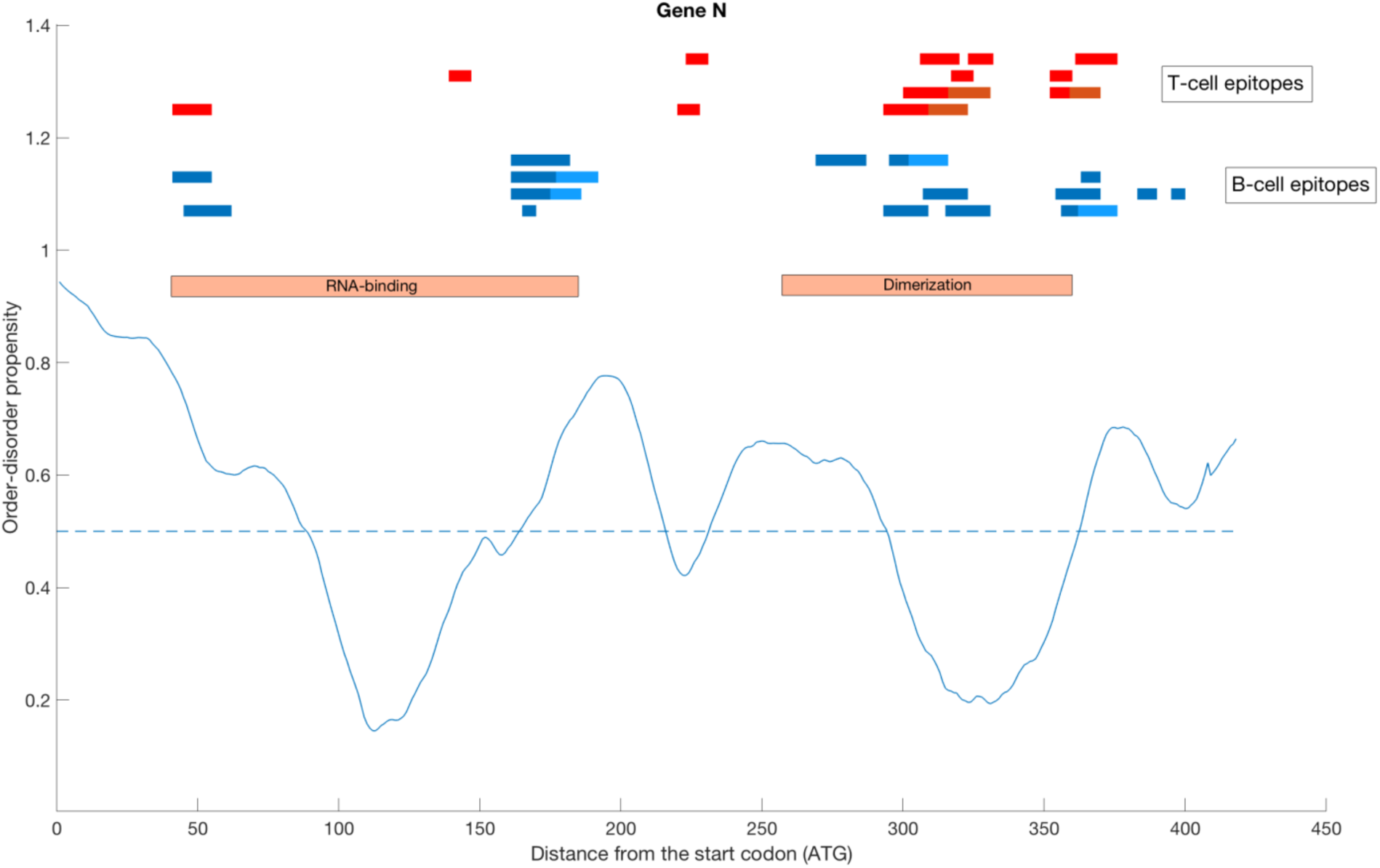
Order-disorder propensity profile of SARS-CoV-2 protein N. The solid blue line represent the position-specific estimations of the structural order–disorder propensity of each residue averaged over a window of 21 residues around that position (taking into account the limitations on both sides of the protein sequence) as a function of the distance from the start codon. Thin horizontal dotted line represent the threshold value, above which the score is characteristic of disorder (0.5 for IUPred2A). On the top, we report the SARS-CoV-2 derived B cell epitopes (in blue) and T cell epitopes (in red) by Ahmed et al. [Ahmed et al. 2020]. With two orange rectangles we show the RNA-binding domain and the dimerization domain.

The rationale to understand the results below is that the score of each residue in the protein sequence ranges from 0 (strong propensity for an ordered structure) to 1 (strong propensity for a disordered structure). In particular, each residue in the sequence is classified as either ordered or disordered depending on whether the IUPred2A score is < 0.5 or > 0.5, respectively. Both the RNA-binding domain and the dimerization domain are predicted to be ordered in a large percentage of their residues, and the vast majority of the B cell epitopes and T cell epitopes overlap these two functional regions of the protein N (see also Fig. S3 for details).

Although disordered protein regions are expected to be under-represented as epitopes due to their sensitive to proteolysis [Mitić et al. 2004], we note the presence of a large fraction of B-cell and T-cell epitopes in predicted disordered regions.

Similarly, we then studied the order-disorder propensity profile of S protein (Fig. 4).

**Figure 4:**
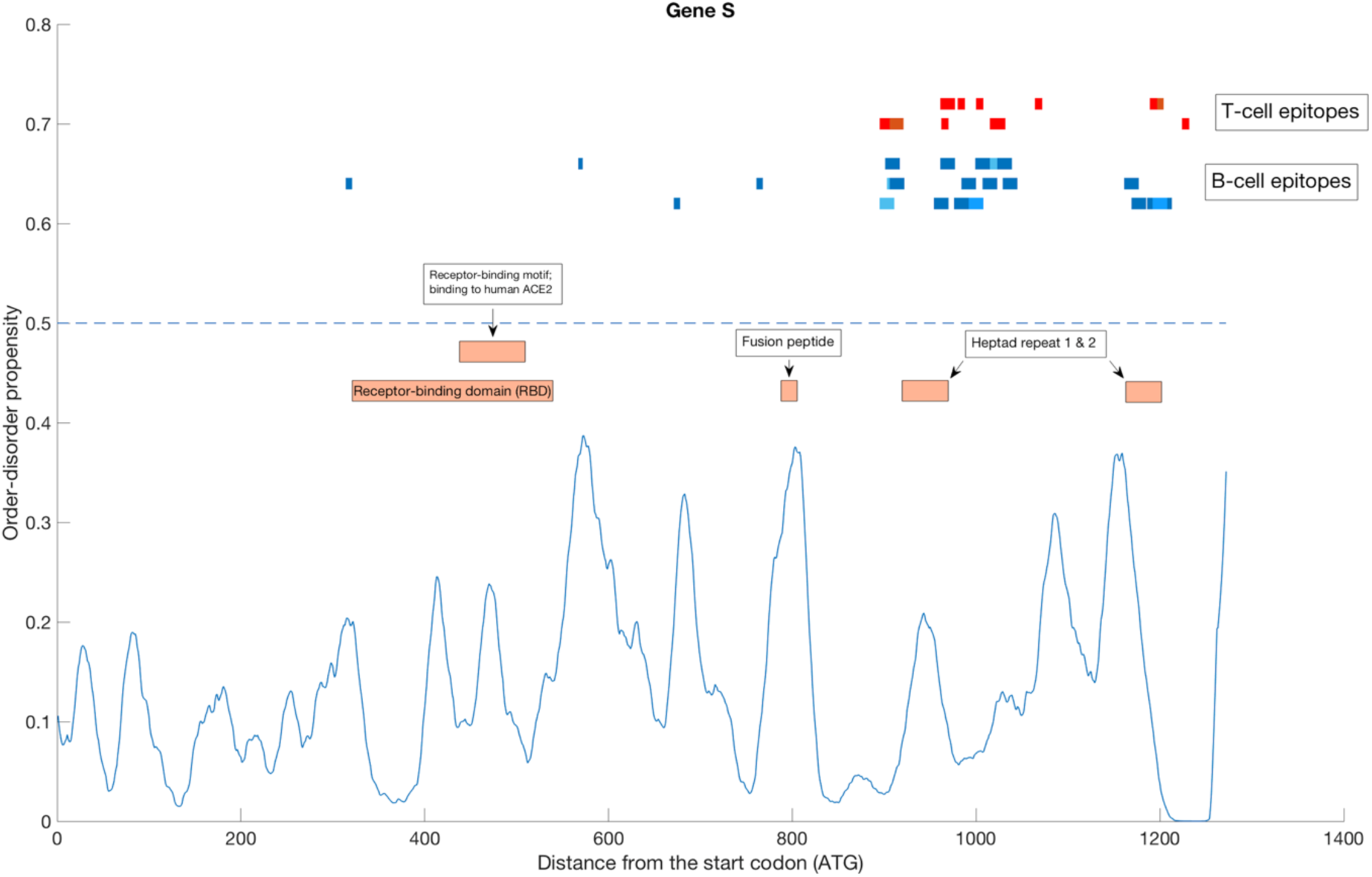
Order-disorder propensity profile of SARS-CoV-2 protein S. The solid blue line represent the position-specific estimations of the structural order–disorder propensity of each residue averaged over a window of 21 residues around that position (taking into account the limitations on both sides of the protein sequence) as a function of the distance from the start codon. Thin horizontal dotted line represent the threshold value, above which the score is characteristic of disorder (0.5 for IUPred2A). On the top, we report the SARS-CoV-2 derived B cell epitopes (in blue) and T cell epitopes (in red) by Ahmed et al. [Ahmed et al. 2020]. With orange rectangles we show the receptor-binding domain and its receptor binding motif to human ACE2, the fusion peptide, and the two heptad repeats 1 and 2.

The spike protein is predicted to be ordered along the whole sequence. Both the receptor binding domain and the fusion peptide are well-structured. Also in this case, we observe that the vast majority of the SARS-CoV-2 derived B cell and T cell epitopes are located in regions displaying reduced disorder tendency (see also Fig. S4 for details).

## 4. Discussion

The characterization of evolutionary and structural properties of the spike-surface (S) glycoprotein and the nucleocapsid (N) protein is central for the development of vaccines and antiviral drugs [Ahmed et al. 2020].

In this study, we performed a systematic analysis of the rates of evolution and the structural order-disorder propensities of protein N and S, in relation to the location of the SARS-CoV-2B cell and T cell epitopes derived from N and S proteins by Ahmed et al. [Ahmed et al. 2020].

Identification of conserved epitopes is of strong interest to help design broad-spectrum vaccines against the present outbreak of SARS-CoV-2. Indeed, high-affinity neutralizing antibodies against conserved epitopes could provide immunity to SARS-CoV-2 and protection against future pandemic viruses. Conservation score measures the evolutionary conservation of an amino acid position in a protein based on the phylogenetic relationships observed amongst homologous sequences. The rate of evolution is not constant among amino acid sites: some positions evolve slowly and are commonly referred to as “conserved”, while others evolve rapidly and are referred to as “variable” (see Fig. 1 and Fig. 2). The rate variations correspond to different levels of purifying selection acting on these sites [Forcelloni and Giansanti 2020]. The purifying selection can be the result of geometrical constraints on the folding of the protein into its 3D structure, constraints at amino acid sites involved in enzymatic activity or in ligand binding or, alternatively, at amino acid sites that take part in protein-protein interactions.

Here, we used Rate4site as a method to infer the rate of evolution of each residue in the amino acid sequences of protein N and S. We then analyzed the structural properties of the protein regions where the epitopes are located by studying their order–disorder propensity.

We show the presence of both conserved epitopes and non-conserved epitopes in terms of rates of evolution. Specifically, the vast majority of the SARS-CoV-2 derived B cell epitopes and T cell epitopes for the nucleocapsid N protein are located in the RNA-binding and dimerization domains. In this case, we found epitopes in both ordered and disordered regions. Although we noted that the vast majority of epitopes are located in regions having high rates of evolution, we also identified epitopes in conserved protein regions. Similarly, we observed that the SARS-CoV-2 derived B cell and T cell epitopes for the spike S protein are preferentially located in protein regions that are predicted to be ordered and conserved. We thus suggest that the immune targeting of these conserved epitopes may potentially offer protection against this novel coronavirus.

Finally, it is worth noting that numerous SARS S-protein-specific neutralizing antibodies have been reported to recognize epitopes within the receptor binding domain (RBD) for ACE2 receptor [Sui et al. 2014]. Indeed, RBD immunization induced specific antibodies may block this recognition and thus effectively prevent the invasion of the virus. However, we showed here that both the receptor-binding motif to ACE2 (region 437-508) and the fusion peptide (region 788-806) are characterized by high rates of evolution, indicating a tendency for these two regions to mutate and, therefore, overcome the host immunity.

In conclusion, our results suggest that targeting conserved regions of SARS-CoV-2 spike and nucleocapsid proteins with less plasticity and more structural constraint should have broader utility for antibody-based immunotherapy, neutralization and prevention of escape variants.

