## Supplementary material for "Identification of conserved epitopes in SARS-CoV-2 spike and nucleocapsid protein": Fig. S1

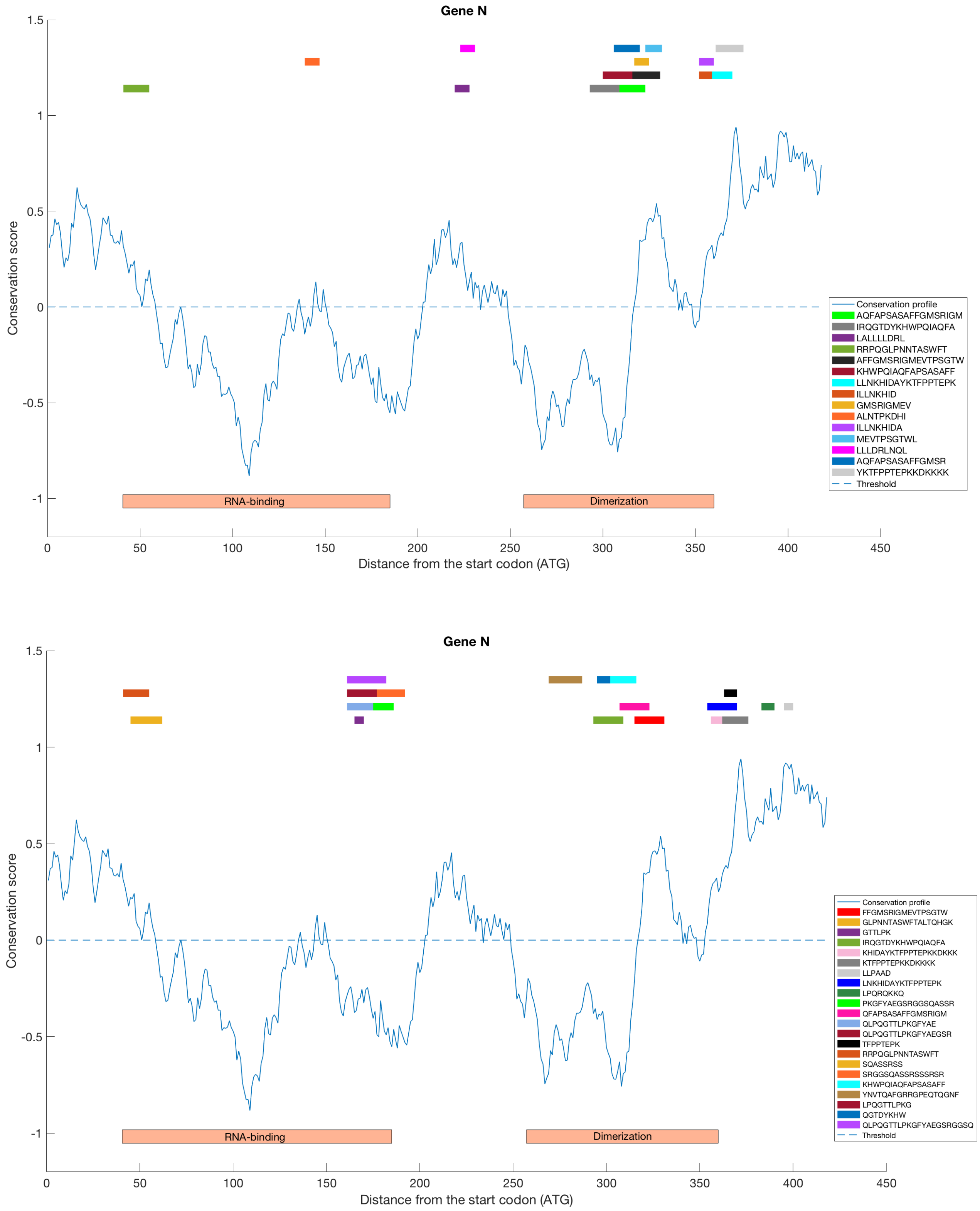


**Figure S1: Conservation profile of SARS-CoV-2 protein N.** The solid blue line represent the position-specific estimations of the rate of evolution of each residue averaged over a window of 11 residues around that position (taking into account the limitations on both sides of the protein sequence) as a function of the distance from the start codon. Thin horizontal dotted line represent the threshold value, above which the score is characteristic of disorder (0 for Rate4site). In each panel, we report the SARS-CoV-2 derived B cell epitopes (on the bottom) and T cell epitopes (on the top) by Ahmed et al. [Ahmed et al. 2020] specifying their sequences in in the legend. With two orange rectangles we show the RNA-binding domain and the dimerization domain.

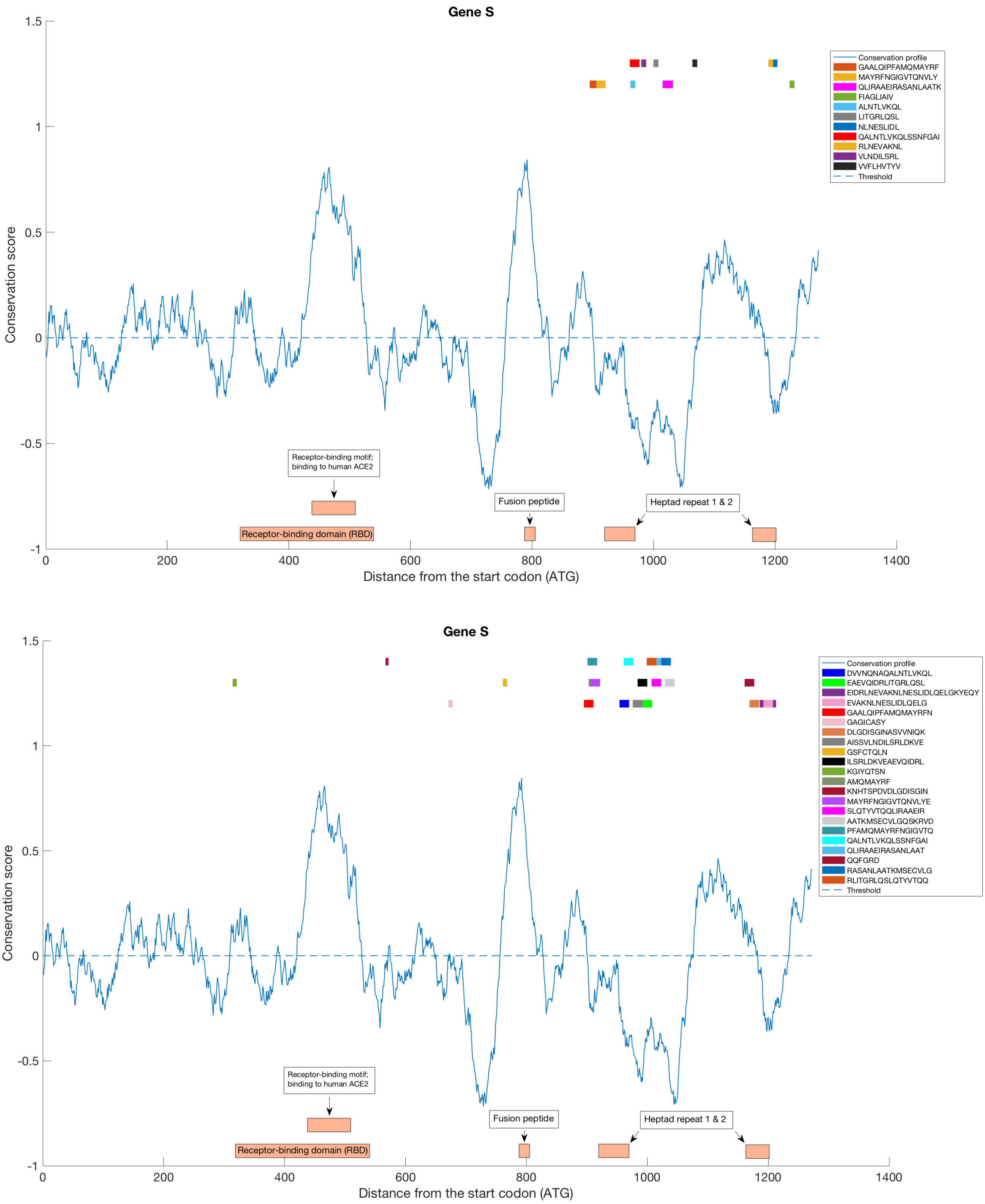


**Figure S2: Conservation profile of SARS-CoV-2 protein S.** The solid blue line represent the position-specific estimations of the rate of evolution of each residue averaged over a window of 11 residues around that position (taking into account the limitations on both sides of the protein sequence) as a function of the distance from the start codon. Thin horizontal dotted line represent the threshold value, above which the score is characteristic of disorder (0 for Rate4site). In each panel, we report the SARS-CoV-2 derived B cell epitopes (on the bottom) and T cell epitopes (on the top) by Ahmed et al. [Ahmed et al. 2020] specifying their sequences in in the legend. With orange rectangles we show the receptor-binding domain and its receptor binding motif to human ACE2, the fusion peptide, and the two heptad repeats.

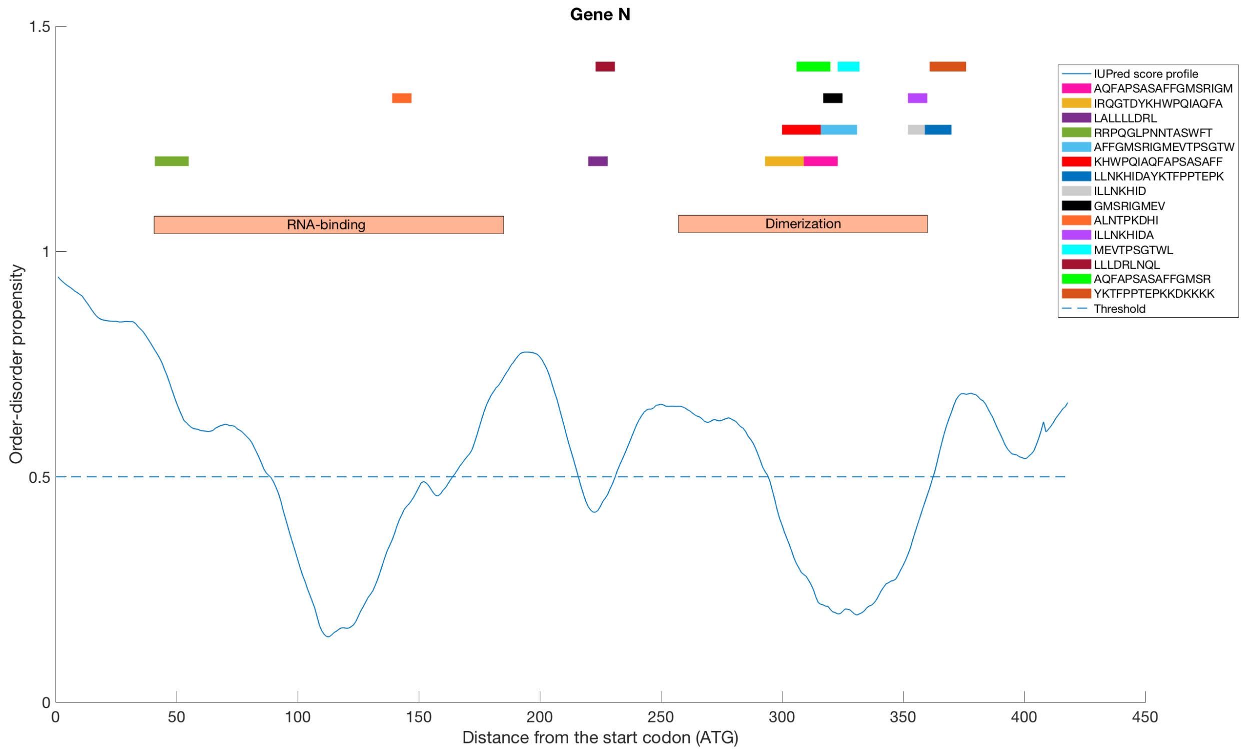


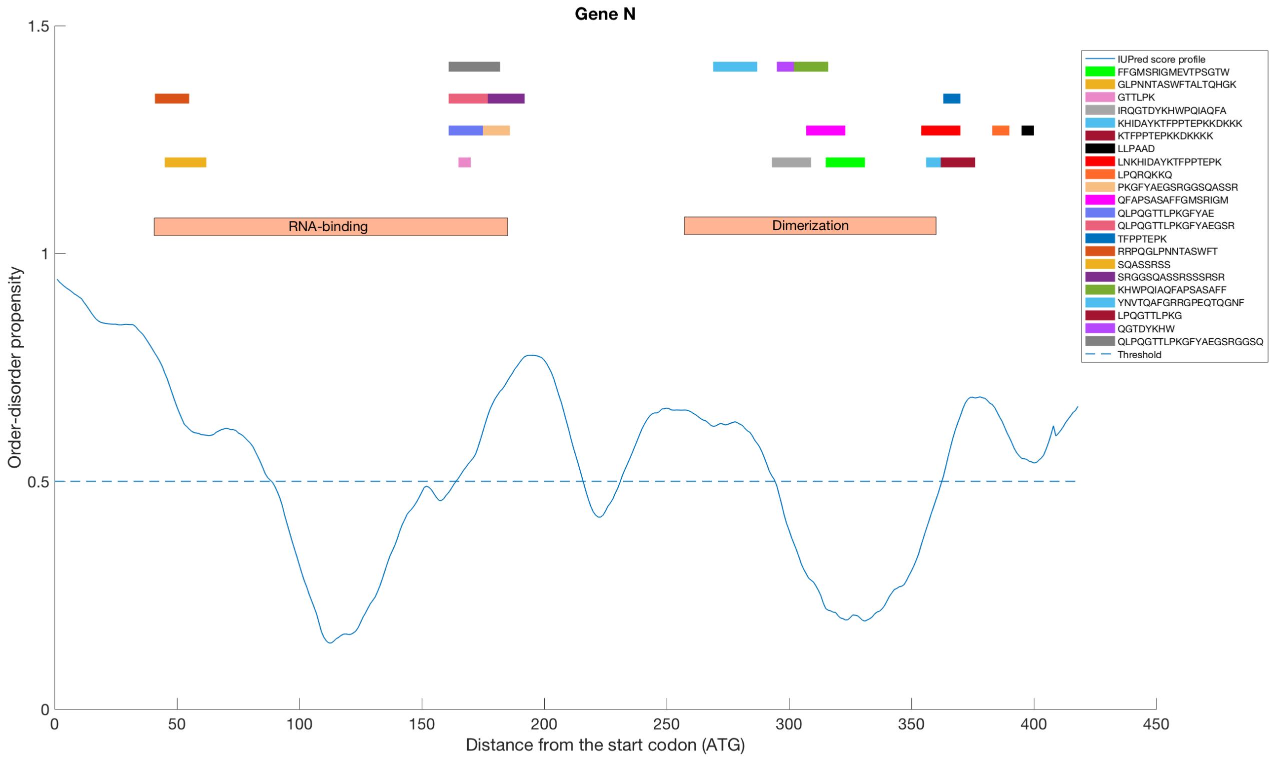


**Figure S3: Disorder profile of SARS-CoV-2 protein N.** The solid blue line represent the position-specific estimations of the structural order–disorder propensity of each residue averaged over a window of 11 residues around that position (taking into account the limitations on both sides of the protein sequence) as a function of the distance from the start codon. Thin horizontal dotted line represent the threshold value, above which the score is characteristic of disorder (0.5 for IUPred2A). In each panel, we report the SARS-CoV-2 derived B cell epitopes (on the bottom) and T cell epitopes (on the top) by Ahmed et al. [Ahmed et al. 2020] specifying their sequences in in the legend. With two orange rectangles we show the RNA-binding domain and the dimerization domain.

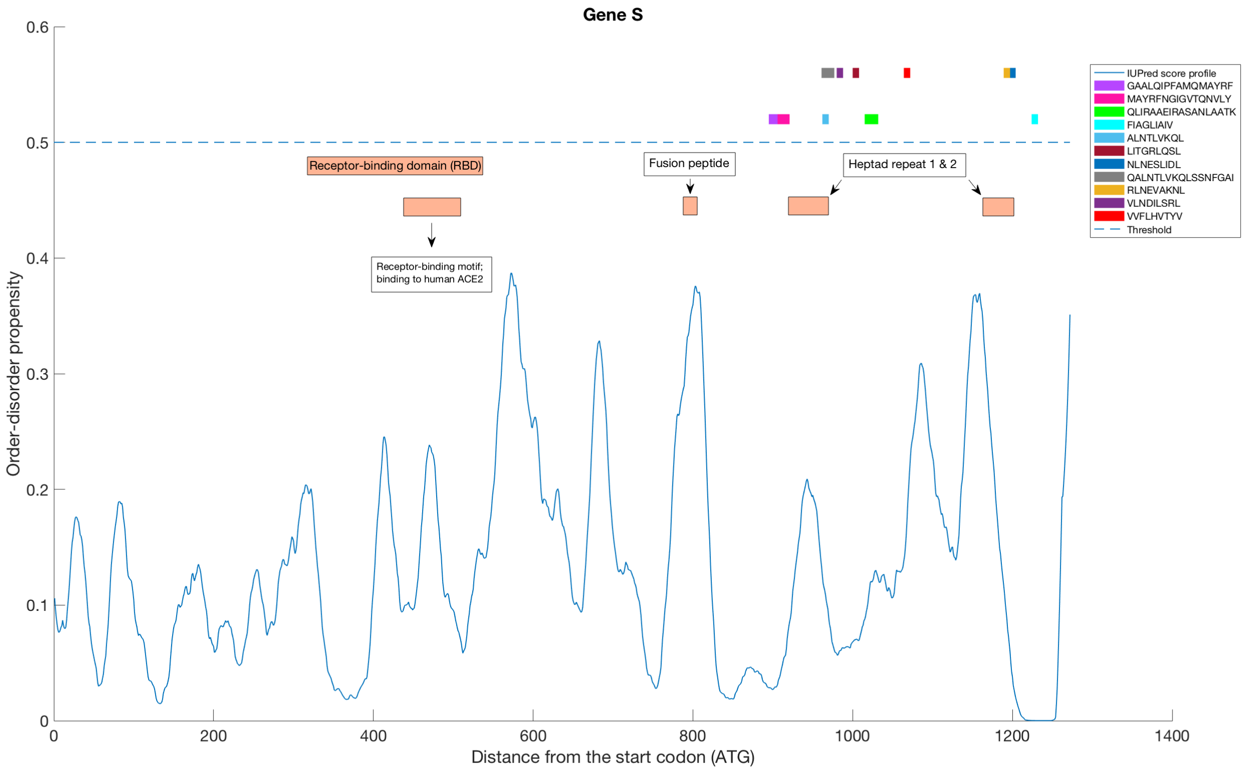


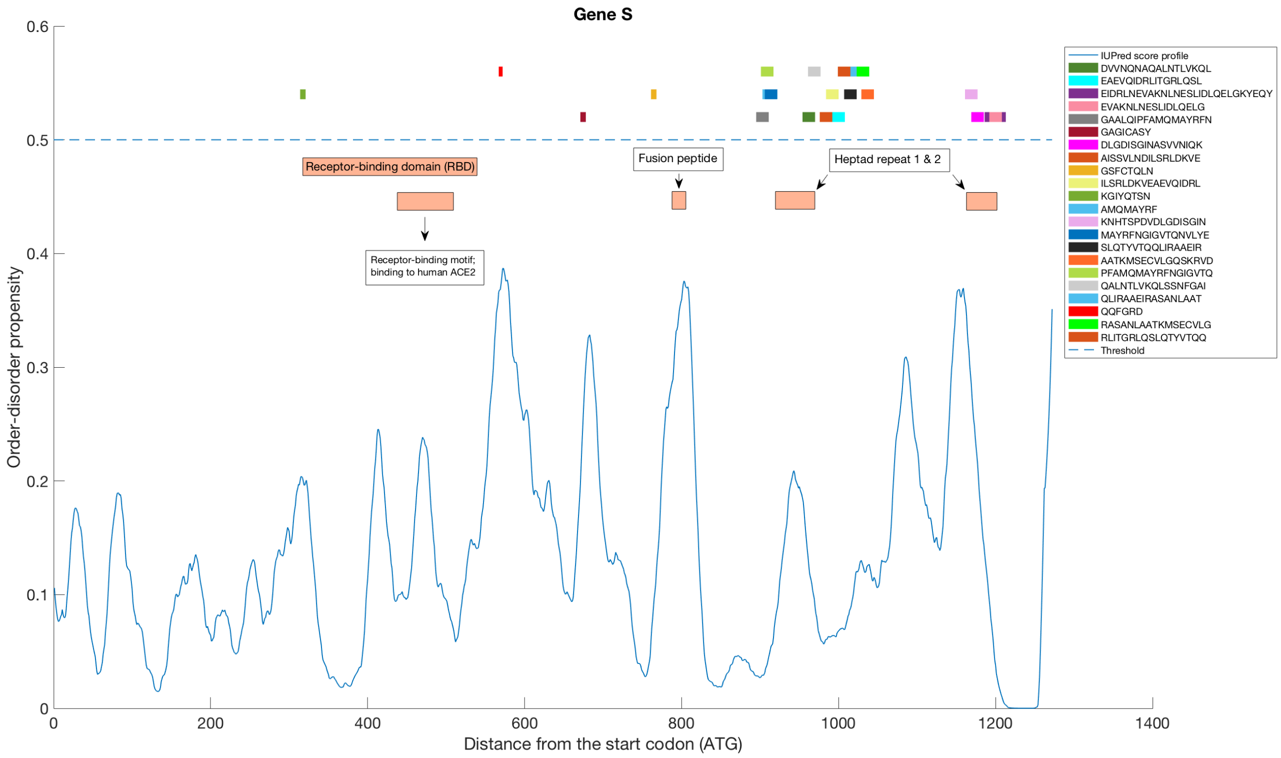


**Figure S4: Disorder profile of SARS-CoV-2 protein S.** The solid blue line represent the position-specific estimations of the structural order–disorder propensity of each residue averaged over a window of 11 residues around that position (taking into account the limitations on both sides of the protein sequence) as a function of the distance from the start codon. Thin horizontal dotted line represent the threshold value, above which the score is characteristic of disorder (0.5 for IUPred2A). In each panel, we report the SARS-CoV-2 derived B cell epitopes (on the bottom) and T cell epitopes (on the top) by Ahmed et al. [Ahmed et al. 2020] specifying their sequences in in the legend. With orange rectangles we show the receptor-binding domain and its receptor binding motif to human ACE2, the fusion peptide, and the two heptad repeats 1 and 2.
